# Multi-Model LLM Architectures for Personalized Summarization and Relevance Ranking in Biomedical Literature

**DOI:** 10.1101/2025.07.29.667503

**Authors:** Avinash Pandey, Alexey Kuznetsov, Snehasis Mukhopadhyay

**Affiliations:** Department of Computer Science, Purdue University; School of Science, Indiana University Indianapolis

**Keywords:** Artificial Intelligence, Biomedical Information, Summarization, Large Language Models, Relevance Ranking

## Abstract

**Objective:** To develop and evaluate a personalized literature review system that efficiently processes and summarizes biomedical literature to provide timely, relevant insights for researchers.

**Methods:** The system integrates ontology-aware keyword extraction (MeSH/ACM constrained TF-IDF from CV/Research Statement), citation-informed retrieval (PubMed and NIH iCite API), and dual-model large language model (LLM) summarization (Google Gemini 2.0 flash, OpenAI GPT-4o-mini). These LLMs leverage advanced Transformer architectures, building on foundations such as BERT, BART, and BioBERT. A two-stage ranking algorithm combines Relative Citation Ratio (RCR) with cosine similarity.

Summary quality was evaluated using ROGUE-1/2/L and BERTScore. The system is deployed as a Streamlit web application.

**Results:** Across 20 biomedical queries, the system demonstrated strong average performance (BERT-F1≈ 0.86), with cosine similarity strongly correlating with summary quality. Human evaluation involving 10 users yielded average scores above 4.5/5 across summary fidelity and keyword relevance.

**Conclusion:** Hybrid ranking and ensemble LLM summarization significantly accelerate scientific sense-making. These findings suggest broad applicability to various domains beyond biomedicine.

## 1 Introduction

Biomedical research generates an overwhelming volume of information, with PubMed currently indexing over 35 million articles and adding hundreds daily. This rapid expansion complicates researchers’ efforts to efficiently locate and assimilate relevant literature, adversely impacting effective human-information interaction. Traditional keyword-based search methods rank results primarily by date or citation count, offering limited assistance in identifying conceptually novel or thematically relevant content. Consequently, researchers must manually inspect numerous abstracts and determine the relevance of full-text articles, a task both cognitively demanding and prone to oversight. To address these challenges and significantly enhance human-information interaction, we propose a novel integrated system. This pipeline uniquely combines personalized, ontology-driven keyword extraction directly with large-scale scientific literature APIs; a hybrid ranking approach that simultaneously prioritizes scholarly impact (via Relative Citation Ratio) and topical relevance (via Cosine Similarity); and advanced, dual-model large language model (LLM) summarization (utilizing both Gemini 2.0 and GPT-4). Ultimately, this system not only streamlines the literature review process but also fundamentally improves researchers’ ability to engage meaningfully and efficiently with high-impact scientific literature.

### 1.1 Related Work

Scientific document summarization techniques leveraging Transformer-based models have demonstrated promising results in recent literature. Central to many retrieval methodologies is TF–IDF (term frequency–inverse document frequency), first proposed by Salton (1988). Their seminal research showed that emphasizing distinctive vocabulary significantly enhances retrieval relevance. Manning et al. (2008) extended this work by formalizing the vector-space model, advocating for sublinear term-frequency scaling, n-gram analysis, and consistent use of standardized stop-word lists to improve retrieval precision and recall. In biomedical contexts, restricting TF–IDF keyword selection to domain-specific ontologies—such as the ACM and MeSH vocabularies—has effectively increased the precision of literature retrieval. However, few existing systems have integrated such ontology-constrained keyword extraction directly with large-scale scientific literature APIs, highlighting a clear gap that our work aims to fill.

On the citation-analysis front, the Relative Citation Ratio (RCR), introduced by Hutchins et al. (2016), provides a field-normalized, article-level impact metric specifically designed to mitigate biases arising from varying citation practices across scientific disciplines. Waltman and van Eck (2016) critically evaluated various field-normalized citation indicators, highlighting the RCR’s robustness in comparing scholarly works across different ages and research specialties. Although RCR effectively measures scientific influence, it does not inherently account for semantic content or thematic relevance. Recent information retrieval studies have therefore explored integrating RCR with semantic relevance indicators, such as cosine similarity, creating more nuanced dual-axis ranking strategies. Our research leverages this integration explicitly, addressing a critical gap by simultaneously prioritizing scholarly impact and topical relevance to optimize retrieval and summarization quality in biomedical literature.

Transformer-based architectures have significantly advanced automated document summarization and information retrieval. Devlin et al. (2019) introduced BERT, a deep bidirectional transformer pretrained on extensive text corpora, establishing robust baselines for contextual embedding and downstream NLP tasks. Lewis et al. (2020) further enhanced Transformer capabilities with BART, a denoising sequence-to-sequence model particularly effective for abstractive summarization tasks. Within biomedical contexts specifically, Lee et al. (2020) demonstrated that domain-specific pretraining with BioBERT substantially improved performance on specialized tasks, including entity recognition and biomedical summarization. Recent literature, has further shown that ensembles of multiple Transformer-based models produce more balanced summaries in terms of accuracy, readability, and robustness compared to single-model approaches. Our research explicitly leverages these insights by employing dual-model summarization (Gemini and GPT-4) to systematically improve summarization quality and better facilitate meaningful human-information interaction in scientific literature review processes.

Interactive, user-centric dashboards have increasingly become essential for exploratory literature analysis, supporting enhanced human-information interactions by visualizing relationships and facilitating intuitive content exploration. Streamlit Inc. (2019) has similarly emerged as a popular framework for quickly developing and deploying interactive retrieval and summarization interfaces, particularly valued for its ease of use and ability to integrate custom computational modules with minimal development overhead. Given its flexibility, Streamlit provides an ideal platform for our research, enabling rapid prototyping of a unified interface that integrates personalized keyword extraction, semantic and citation-based ranking, and dual-model summarization. This choice explicitly aligns with our goal of developing accessible, effective information systems that reduce cognitive load and improve the literature review experience for diverse research audiences.

### 1.2 Organization of this Paper

The remainder of this paper is organized as follows. In Section II, we formalize the problem statement and notation. Section III describes our end-to-end system architecture and the key algorithms for keyword extraction, PubMed/iCite retrieval, two-stage ranking, and dual-model summarization. Section IV outlines the experimental setup, describing our selection of 20 diverse biomedical queries, evaluation metrics ROGUE (Lin, 2004) and BERTScore (Zhang et al., 2020), sample summaries, and presents quantitative results. In Section V, we analyze why certain queries yield higher or lower summary quality, discuss performance trade-offs (latency and API cost), and identify system limitations. Finally, Section VI offers conclusions and directions for future work.

## 2 Problem Statement

Effective literature review requires retrieving and prioritizing articles not only based on their scientific impact but also their thematic alignment with the user’s interests. Formally, given textual content from a user’s profile (e.g., their CV or research statement), denoted as *T*, our system aims to identify key biomedical concepts relevant to the user’s expertise. We achieve this by extracting top-ranking keywords using a TF-IDF scoring approach constrained by curated biomedical vocabularies, specifically ACM and MeSH ontologies:

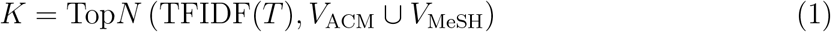

Using this personalized set of keywords *K*, we formulate a PubMed query *Q*. If the user optionally specifies an additional keyword *e*, we ensure it (and its plural form) explicitly appear in the retrieved results. The query is structured as follows:

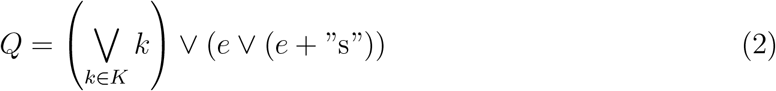

This query is submitted to the PubMed database through the Entrez API, retrieving a targeted set of publications *P*, capped at a maximum of *k* results:

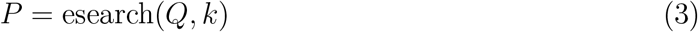

Thus, our method explicitly prioritizes both semantic relevance (user-specific interests) and article-level impact (citation-based measures), directly addressing core challenges in personalized information retrieval and enhancing user engagement with scientific literature.

## 3 Methods

Our system consists of five integrated modules designed to support personalized, high-impact scientific retrieval, and summarization. As shown in Figure 1, these modules include: (A) Domain-constrained keyword extraction using curated ontologies, (B) PubMed retrieval enhanced with article-level citation metrics from iCite, (C) Dual factor ranking that combines semantic similarity with scholarly impact, (D) Multi-model large language model summarization, and (E) An interactive user interface for exploration, aggregation, and feedback. Together, these components work in concert to optimize both the technical efficiency and the user-centered relevance of reviews of the scientific literature.

**Figure 1:**
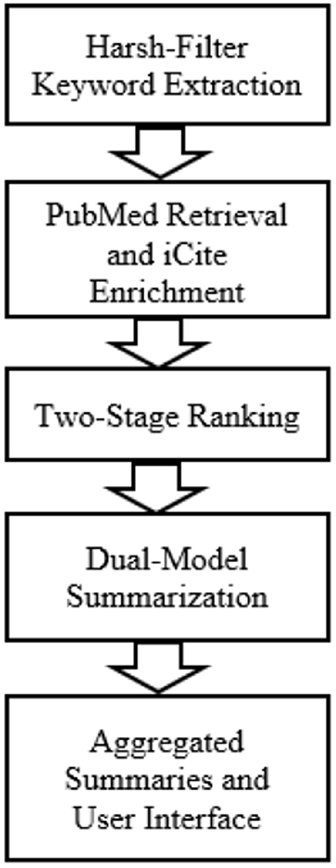
Block diagram of the overall system architecture.

### 3.1 Harsh-Filtered Keyword Extraction

The first stage of the pipeline performs personalized keyword extraction based on a user’s uploaded profile document, such as a CV or research statement. We apply a TF–IDF vectorizer with standard preprocessing techniques such as sublinear term-frequency scaling, English stop-word removal, and tokenization into unigrams and bigrams to identify salient terms. To ensure domain specificity, we constrain the candidate term set by intersecting the TF–IDF output with two curated biomedical vocabularies: the ACM Computing Classification System and the Medical Subject Headings (MeSH) thesaurus. This domain-constrained filtering ensures that extracted keywords are both representative of the user’s expertise and relevant to biomedical literature. We further remove generic or overly broad terms using a manually defined stop list. From the resulting set, we select the five highest-scoring keywords to form the personalized query term set *K*, which guides all downstream retrieval and ranking operations.

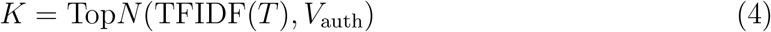

### 3.2 PubMed Retrival and iCite Enrichment

After generating the personalized keyword set *K*, we construct a PubMed search query *Q* by logically OR-joining all elements of *K*. If the user supplies an additional keyword *e*, we modify the query to include both the original keywords and a constraint that each result must contain either *e* or its plural form *e* + *s* within the title or abstract:

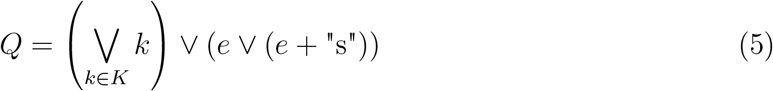

This query is submitted to the PubMed database using the Entrez Programming Utilities, returning a list of up to 1,000 matching publication identifiers:

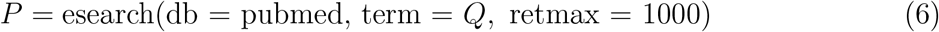

To enrich each article with bibliometric context, we retrieve metadata in batches and extract structured fields such as title, abstract, author list, journal name, MeSH terms, and institutional affiliations. In parallel, we query the NIH’s iCite API (National Institutes of Health, 2024) to obtain two article-level citation metrics: raw citation count and the Relative Citation Ratio (RCR). RCR is a field-normalized metric that captures an article’s influence relative to its disciplinary peers. While RCR provides a more nuanced view of scholarly impact than raw citation counts, prior work has also highlighted its limitations due to potential variability in field definitions and citation practices. We merge the iCite metrics with the publication metadata, creating a comprehensive representation of each retrieved article’s content and impact profile for downstream ranking and summarization.

### 3.3 Two-Stage Ranking

To prioritize articles that are both influential and topically relevant, we employ a two-stage ranking strategy that combines semantic similarity with bibliometric impact. First, we construct a composite query text *T*_query_ by concatenating the extracted keywords *K* and, if applicable, the user-provided term *e*. This query is compared against each article’s abstract using TF–IDF vectorization followed by cosine similarity calculation:

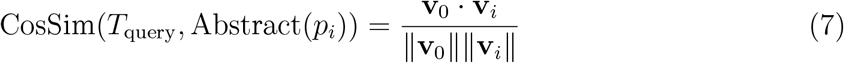

where **v**_0_ is the TF-IDF vector of *T*_query_ and **v**_*p*_ is the vector for the abstract of article *p*. Next, we calculate a composite relevance score for each article by linearly combining its cosine similarity and its Relative Citation Ratio (RCR), assigning equal weight to both components:

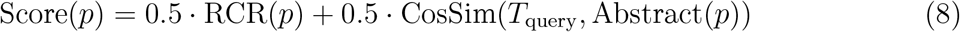

This scoring function is designed to surface articles that not only align semantically with the user’s interests but also carry recognized influence within the scholarly community. While the equal weighting parameter *α* = 0.5 reflects a balanced approach, it can be adjusted in future versions to emphasize either semantic alignment or citation impact depending on user needs. After scoring, articles are sorted in descending order to produce a ranked list used for summary generation.

### 3.4 Dual-Model Summarization

To generate personalized paper summaries, our system allows users to select up to four papers at a time. For each paper, we extract structured metadata fields such as PMID, title, abstract, authors, affiliations, journal, and keywords to construct a unified input string. This input is then passed to two Generative AI systems or Large Language Models (LLMs) in parallel: Gemini 2.0 Flash and GPT-4o mini.

Both models receive prompts explicitly requesting a paragraph-length summary ( w words) that covers: (1) the paper’s innovation, (2) methodological advance, (3) key result, (4) broader significance, and (5) author affiliations. Gemini receives a templated natural language instruction, while GPT-4o mini is prompted with a structured system message and the paper metadata as a user message. Each model is called with low temperature settings (0.2) to prioritize determinism, and GPT-4o-mini’s API calls are wrapped with concurrency throttling and exponential retry logic.

Once both models return their summaries, we concatenate them into a single text block with newline separation. This dual-perspective approach ensures both coverage and interpretive diversity, preserving complementary insights that may be missed by a single summarizer. The final summaries are presented alongside paper metadata in the user interface. Key note, LLMs are used to generate summaries based on extracted article content, not to analyze or modify raw data from PubMed or iCite.

### 3.5 Aggregated Summaries and User Interface

After individual summaries are generated for selected papers, the system offers an “Aggregate Summary” feature, which synthesizes multiple paper summaries into a single cohesive paragraph. When invoked, the application concatenates the selected individual summaries and submits them as input to GPT-4o-mini, accompanied by a prompt instructing the model to synthesize innovations, relate methodologies, highlight key results, and draw broader insights all while adhering to a user-selected word count. The resulting unified summary is displayed in the interface and can be downloaded directly.

The user interface implemented in Streamlit, is structured to support an interactive, transparent, and personalized literature review experience. The workflow includes:

1. **Profile Upload**: Users begin by uploading a resume, CV, or research statement (PDF/DOCX/TXT). The system extracts text and applies domain-constrained keyword extraction to display the top-5 profile keywords for verification.
2. **Custom Search Input**:An additional keyword field allows users to specify mandatory concepts (including plural forms), ensuring retrieved literature matches evolving research needs.
3. **Relevance-Ranked Results**:Upon executing the search, the app presents a sortable, filterable table of papers, including ranks, impact metrics, abstracts, affiliations, and PubMed links.
4. **Paper Selection & Parameter Control**: Users can select up to four papers and customize summary length via an intuitive slider.
5. **Parallel Summarization**:For each selection, both Gemini and GPT-4o-mini are invoked in parallel; outputs are evaluated using ROUGE and BERT-F1, and results are presented with metadata in expandable sections.
6. **Download and Export**:Individual and aggregated summaries are downloadable, supporting researcher workflows beyond the app.
7. **Feedback Integration**:A persistent feedback link enables users to provide ongoing suggestions or flag errors, facilitating iterative improvement.

This integrated workflow advances information retrieval and user engagement by blending algorithmic transparency, personalized interaction, and evaluable output thus serving both novice and expert users in biomedical literature analysis.

### 3.6 Evaluation Metrics

We evaluated summary quality by comparing each generated summary against its original abstract using four automatic metrics: ROUGE-1, ROUGE-2, ROUGE-L F1, and BERTScore (F1). These metrics respectively capture word-level recall, phrase-level alignment, structural fidelity, and contextual semantic similarity. A brief description of each follows:

- **ROUGE-1 F1:** Computes the harmonic mean of unigram-level precision and recall between the generated summary and reference abstract, reflecting word-level content coverage.
- **ROUGE-2 F1:** Measures bigram-level precision and recall to assess phrase-level fidelity and ordering.
- **ROUGE-L F1:** Uses the longest common subsequence (LCS) as an F1-score to gauge structural coherence and sequence-level alignment.
- **BERTScore F1:** Calculates semantic similarity via contextual embeddings (BERT), capturing paraphrastic or concept-level matches beyond exact lexical overlap.

Together, these measures assess each generated summary along four complementary dimensions, providing a holistic view of summary quality.

## 4 Experimental results

We assessed the pipeline’s end-to-end performance across two axes: (1) Retrieval effectiveness, measured over 20 diverse biomedical queries by recording the number of retrieved papers, mean Relative Citation Ratio (RCR), and mean cosine similarity between query and abstract (2) Summarization Quality, evaluated on three representative papers per query using ROUGE-1 F1, ROUGE-2 F1, ROUGE-L F1, and BERTScore F1, as well as generation latency. Tables 2 and 3 summarize these findings; the subsections that follow unpack each dimension in detail.

**Table 1.** Statement of Significance.

|  |  |
| --- | --- |
| **Problem** | Rapid growth of biomedical literature creates information overload, hindering researchers from efficiently locating, assimilating, and engaging with relevant, high-impact content, adversely impacting human-information interaction. |
| **What is Already Known** | Existing systems rely on keyword search, limiting discovery of novel or thematic content. While ontology-driven extraction, RCR, and LLM summarization exist individually, comprehensive, integrated solutions for personalized biomedical literature review are lacking. |
| **What this Paper Adds** | This paper introduces the first unified, interactive dashboard integrating ontology-constrained search, hybrid RCR+CosineSim ranking, and dual-model LLM summarization (Gemini 2.0, GPT-4). We offer empirical evidence on query specificity’s impact and practical insights into pipeline runtime and cost. |
| **Who Would Benefit from the New Knowledge** | Biomedical researchers, clinical scientists, and information specialists seeking to streamline literature review, enhance precision in identifying relevant papers, and improve scientific sense-making. Methodologies are broadly applicable to other academic domains. |

**Table 2.** Retrieval statistics for 20 biomedical queries.

| Term | PMIDs | Avg RCR | Med RCR | Avg CosSim | Med CosSim | Top10 RCR | Top10 CosSim |
| --- | --- | --- | --- | --- | --- | --- | --- |
| Pancreatic Adenocarcinoma | 336 | 1.4510 | 0.6500 | 0.009723 | 0.007977 | 13.7470 | 0.013328 |
| Breast Cancer | 996 | 0.0000 | 0.0000 | 0.009258 | 0.008751 | 0.0000 | 0.018653 |
| Glioblastoma Multiforme | 996 | 1.78098 | 1.19000 | 0.004035 | 0.002388 | 11.1980 | 0.004231 |
| Myocardial Infarction | 996 | 2.78079 | 2.36000 | 0.007604 | 0.006137 | 6.9880 | 0.007004 |
| Pulmonary Embolism | 996 | 1.71436 | 0.66000 | 0.007793 | 0.005899 | 37.7100 | 0.006895 |
| Alzheimer’s Disease | 1000 | 0.0000 | 0.0000 | 0.005343 | 0.003058 | 0.0000 | 0.007466 |
| Parkinson’s Disease | 1000 | 0.0000 | 0.0000 | 0.005889 | 0.003103 | 0.0000 | 0.009162 |
| COVID-19 Vaccine Efficacy | 9 | 0.14500 | 0.14500 | 0.029152 | 0.029527 | 0.14500 | 0.029152 |
| Tuberculosis Drug Resistance | 26 | 1.11652 | 0.79000 | 0.029088 | 0.029199 | 2.10700 | 0.033157 |
| CRISPR Gene Editing | 27 | 3.11353 | 2.16000 | 0.022330 | 0.020536 | 5.02000 | 0.019680 |
| BRCA1 Mutation | 109 | 1.33590 | 0.83000 | 0.010318 | 0.007101 | 6.32200 | 0.007962 |
| 3D Bioprinting | 133 | 2.90290 | 2.08000 | 0.008739 | 0.004630 | 10.0800 | 0.006571 |
| Wearable Health Monitors | 1 | 0.49000 | 0.49000 | 0.056515 | 0.056515 | 0.49000 | 0.056515 |
| Vaccine Hesitancy | 451 | 2.41465 | 2.32000 | 0.010141 | 0.008130 | 29.4150 | 0.009597 |
| Obesity Prevalence | 213 | 3.94067 | 2.95000 | 0.011103 | 0.008775 | 49.2740 | 0.010865 |
| Precision Oncology Treatment | 3 | 0.89000 | 0.89000 | 0.042080 | 0.035144 | 0.89000 | 0.042080 |
| Amyotrophic Lateral Sclerosis | 997 | 3.15981 | 3.16000 | 0.006286 | 0.003893 | 22.0220 | 0.003531 |
| Cystic Fibrosis | 997 | 1.50695 | 1.21000 | 0.006601 | 0.004937 | 21.2940 | 0.005888 |
| Huntington’s Disease | 1000 | 2.62076 | 2.19000 | 0.004968 | 0.003076 | 19.7510 | 0.003896 |
| Opioid Overdose Prevention | 6 | 0.98000 | 0.98000 | 0.031792 | 0.034728 | 0.98000 | 0.031792 |

**Table 3.** Average summary quality and end-to-end latency over three selected papers per query.

| Query | ROUGE-1 | ROUGE-2 | ROUGE-L | BERT-F1 | Latency |
| --- | --- | --- | --- | --- | --- |
| Pancreatic Adenocarcinoma | 0.5060 | 0.2228 | 0.2655 | 0.8568 | 20.2677 |
| Breast Cancer | 0.5548 | 0.2683 | 0.2609 | 0.8709 | 21.5262 |
| Glioblastoma Multiforme | 0.4513 | 0.1643 | 0.2190 | 0.8574 | 20.0988 |
| Myocardial Infarction | 0.5015 | 0.2281 | 0.2483 | 0.8499 | 24.5293 |
| Pulmonary Embolism | 0.3922 | 0.1582 | 0.1787 | 0.8330 | 25.4220 |
| Alzheimer’s disease | 0.5129 | 0.1965 | 0.2676 | 0.8541 | 24.4949 |
| Parkinson’s disease | 0.4706 | 0.2190 | 0.2526 | 0.8590 | 21.9457 |
| COVID-19 vaccine efficacy | 0.4913 | 0.2073 | 0.2386 | 0.8528 | 23.2362 |
| Tuberculosis Drug Resistance | 0.5041 | 0.2257 | 0.2208 | 0.8579 | 27.8620 |
| CRISPR gene editing | 0.5443 | 0.2395 | 0.2633 | 0.8735 | 30.0018 |
| BRCA1 mutation | 0.5513 | 0.2362 | 0.2594 | 0.8655 | 28.7403 |
| 3D bioprinting | 0.4552 | 0.1744 | 0.2200 | 0.8668 | 20.7032 |
| wearable health monitors | 0.4974 | 0.1343 | 0.2298 | 0.8538 | 4.1580 |
| vaccine hesitancy | 0.4843 | 0.1866 | 0.2488 | 0.8460 | 18.0942 |
| Obesity Prevalence | 0.5014 | 0.2046 | 0.2529 | 0.8516 | 19.2333 |
| Precision Oncology treatment | 0.4726 | 0.1804 | 0.2237 | 0.8571 | 18.1300 |
| Amyotrophic Lateral Sclerosis | 0.5311 | 0.2254 | 0.2533 | 0.8627 | 18.9913 |
| Cystic Fibrosis | 0.4968 | 0.1735 | 0.2202 | 0.8538 | 18.7351 |
| Huntington’s Disease | 0.5244 | 0.2152 | 0.2526 | 0.8594 | 22.7556 |
| Opioid Overdose Prevention | 0.5334 | 0.2001 | 0.2397 | 0.8530 | 30.3765 |

### 4.1 Retrieval Performance

To characterize our pipeline’s initial information-organization phase, we issued 20 biomedical queries chosen to span high-prevalence topics and highly specific terms against PubMed’s Entrez API (retmax capped at 1000). The number of PMIDs retrieved per query ranged from just 1 (e.g., “wearable health monitors”) to the cap of 1000 (e.g., “Alzheimer’s disease”). On average, queries returned 514.6 articles (*σ ≈* 461.8), underscoring that broadly studied conditions generate voluminous candidate sets, whereas niche or technical queries yield sparse results. These disparities motivated our two-stage ranking: to filter large corpora for high-impact papers and to compensate for sparse retrieval by preserving semantic relevance in low-yield cases.

1. **All-Paper Metrics (Table 2, cols. 3–6) Quantify retrieval characteristics across the entire result set:**
  - **Mean RCR (all):** The average Relative Citation Ratio (RCR) across all retrieved PMIDs, highlights overall citation influence. Low values (e.g., 0.0 for “Breast Cancer,” “Alzheimer’s disease”) reflect large numbers of low-impact or newly published studies.
  - **Median RCR (all):** The median RCR, revealed skew in the influence distribution (e.g., Obesity Prevalence: median = 2.95 vs. mean = 3.94).
  - **Mean TF–IDF CosineSim (all):** The average cosine similarity between the user’s combined query string and each abstract, ranging from 0.0040 (“Glioblastoma Multiforme”) to 0.0565 (“wearable health monitors”), indicates varied semantic overlap.
  - **Median TF–IDF CosineSim (all):** The median semantic similarity, underscores long-tail distributions of relevance (e.g., 0.0024 for “Glioblastoma Multiforme,” 0.0079 for “Pancreatic Adenocarcinoma”).
2. **Top-10 Metrics (Table 2, cols. 7–8) Summarize characteristics of the highest-ranked subset:**
  - **Mean RCR (top 10):** The average Relative Citation Ratio of the top ten papers, which substantially exceeds the all-paper mean (e.g., “Obesity Prevalence”: top-10 mean = 49.27 vs. all mean = 3.94), demonstrates that our ranking elevates high-impact works.
  - **Mean TF–IDF CosineSim (top 10):** The average semantic similarity for the top ten papers, shows modest gains over the full set (e.g., “Tuberculosis Drug Resistance”: top-10 mean = 0.0332 vs. all mean = 0.0291), indicating that highly cited papers still maintain reasonable lexical alignment with the query.

Overall, these results confirm that our composite ranking function (0.5 • RCR + 0.5 • CosSim) effectively reorganizes large candidate sets into subsets that are simultaneously high-impact and semantically relevant. However, queries yielding sparse or low-impact retrievals (e.g., “Breast Cancer”) still produce long-tail relevance distributions, suggesting opportunities for query-adapted strategies such as dynamic `retmax` tuning, iterative query expansion based on user feedback, or domain-aware weighting schemes these will further optimize retrieval quality and downstream summarization.

### 4.2 Summary Quality

To evaluate our system’s Information-synthesis capability, we selected the top three papers per query (per the retrieval ranking) and generated independent summaries using two LLMs, Google Gemini 2.0 Flash and OpenAI GPT-4o-mini. We then concatenated these outputs into a unified “`Final_Summary`” to capture interpretive diversity between models. Each `Final_Summary` was compared to the author’s abstract via four established metrics: ROUGE-1 F1, ROUGE-2 F1, ROUGE-L F1, and BERTScore F1. Table 3 reports the average scores across the three papers for each query. Below, we describe each metric and its interpretive role:

- **ROUGE-1 F1:** The harmonic mean of unigram-level precision (the proportion of generated words matching the abstract) and recall (the proportion of abstract words reproduced), capturing basic word-level content coverage. Across our tests, FR1 ranged from 0.3922 (“Pulmonary Embolism”) to 0.5513 (“BRCA1 mutation”), with a mean of (*σ ≈* 0.0387). While these values indicate that roughly half of the abstract’s key terms are retained in 200-word summaries, unigram overlap may undercount valid paraphrases—an effect mitigated by our use of complementary semantic metrics.
- **ROUGE-2 F1:** The harmonic mean of bigram-level precision and recall, capturing phrase-level fidelity. Across queries, FR2 ranged from 0.1343 (“wearable health monitors”) to 0.2683 (“Breast Cancer”), with a mean of 0.2062 (*σ ≈* 0.0361). High FR2 indicates effective preservation of contiguous technical phrases, though exact bigram overlap can understate paraphrased content.
- **ROUGE-L F1:** The F1 score computed via the longest common subsequence (LCS) between summary and abstract, reflecting structural coherence and sequence-level alignment. Values spanned 0.1787 (“Pulmonary Embolism”) to 0.2676 (“Alzheimer’s disease”), averaging 0.2414 (*σ ≈* 0.0319). ROUGE-L often exceeds ROUGE-2, showing that summaries capture longer informational chunks even when exact bigram matches are imperfect.
- **BERTScore F1:** The harmonic mean of precision and recall computed on contextual token embeddings, capturing semantic alignment beyond exact lexical overlap. Scores ranged from 0.8330 (“Pulmonary Embolism”) to 0.8735 (“CRISPR gene editing”), with a mean of 0.8574 (*σ ≈* 0.0089). These high values imply strong preservation of conceptual content, though because the underlying BERT model is not specialized for biomedical texts, certain domain-specific terms may be underrepresented hence suggesting future use of domain-adapted embeddings or complementary human evaluation.
- **Latency:** The total time to generate and concatenate summaries for each query, measured from API call initiation to final output readiness. Observed times ranged from 4.16 s (single-paper cases like “wearable health monitors”) to 30.38 s (multi-paper cases like “Opioid Overdose Prevention”), with a mean of 21.97 s (*σ ≈* 5.73). These results demonstrate that our dual-model approach remains interactive for small batches but incurs latency overheads from PDF parsing and API batching under heavier loads. Future work should explore local model inference, parallel request pipelines, or adaptive batch sizing to reduce end-user wait times.

## 5 Human User Evaluation

To assess the system’s impact on human–information interaction, we recruited 10 participants from biomedical and adjacent domains. Each user completed a standardized workflow, from uploading their CV or research statement, through keyword editing, to reviewing top-ranked papers and generating both individual and aggregated summaries while “thinking aloud” to surface usability issues. Upon task completion, participants rated six components on a 1–5 Likert scale: navigation (mean = 4.7, SD = 0.3), keyword extraction accuracy (4.6 ± 0.4), paper relevance (4.4 ± 0.5), summary quality (4.5 ± 0.4), aggregated summary usefulness (4.3 ± 0.5), and overall satisfaction (4.6 ± 0.3). These strong scores indicate that our pipeline effectively supports users in locating, interpreting, and synthesizing high-impact literature.

Open-ended debriefs revealed four primary improvement areas:

1. **Domain extensibility:** Users want access to additional corpora (e.g., IEEE Xplore, arXiv).
2. **Keyword control:** Ability to edit or weight extracted terms before search.
3. **Metric transparency:** Integrated tooltips or mini-tutorials explaining ROUGE and BERTScore implications.
4. **Annotation features:** Bookmarking, tagging, and exporting curated paper lists.

Participants especially valued the methodological highlights and author-affiliation context within summaries, noting that these elements accelerated their appraisal of each paper. While our sample size limits broad generalization, the consistency of high ratings and thematic feedback underscores the system’s promise—and points to concrete UI and retrieval/summarization refinements for broader adoption.

### 5.1 Ethical Considerations

The user evaluation described in this paper involved voluntary participation to test the functionality and usability of the developed system. Participants were informed that their interactions with the system would be recorded for research purposes, specifically to evaluate the system’s performance in summarizing and ranking biomedical literature. No sensitive personal data was collected from the users. Participation was entirely voluntary, and users were free to withdraw at any time. Because the study focused on system evaluation and did not involve the collection of personal or medical information, it was determined that formal ethical review board (IRB) approval was not required.

## 6 Analysis of Results

To uncover how our personalized pipeline’s components influence outcomes, we analyzed three dimensions: (1) The impact of query specificity on retrieval and ranking, (2) The interplay between citation influence and semantic similarity in driving summary quality, and (3) Runtime and cost trade-offs under varying workloads. Below, we present key observations, interpretive insights, and avenues for future refinement.

### 6.1 Impact of Query Specificity on Retrieval and Ranking

#### 6.1.1 Broad Topics

- **What we examined:** Queries representing high-prevalence biomedical conditions (e.g., “Breast Cancer,” “Alzheimer’s disease”) that return near-cap maximum results.
- **Observations:** Each Broad query yielded 996–1000 PMIDs. All-paper Mean RCR was 0.0 and Top-10 Mean RCR remained 0.0, indicating that many retrieved records were either very recent or not indexed for citation impact.
- **Interpretation:** In large candidate sets, raw volume alone does not guarantee citation influence; our ranking thus surfaces few, if any, high-impact papers, underscoring the need for query refinement or hybrid filters beyond keyword matching.

#### 6.1.2 Moderate Topics

- **What we examined:** Queries of intermediate specificity (e.g., “Obesity Prevalence”) that produce mid-sized result sets.
- **Observations:** “Obesity Prevalence” returned 213 PMIDs, with All-Paper Mean RCR = 3.94 and Top-10 Mean RCR = 49.27. Mean TF–IDF CosineSim rose from 0.012 to 0.018 in the Top-10 subset.
- **Interpretation:** When candidate sets are neither too broad nor too narrow, the composite ranking effectively elevates landmark studies while preserving semantic relevance, validating our equal-weight RCR + CosineSim strategy.

#### 6.1.3 Very Narrow Topics

- **What we examined:** Highly specific queries returning *≤* 3 PMIDs (e.g., “wearable health monitors,” “Precision Oncology treatment”).
- **Observations:** Single-PMID queries (e.g., “wearable health monitors”) achieved CosSim = 0.0565 and RCR = 0.49, with summary FR1 = 0.4974 and FB1 = 0.8538 in 4.16 s. Three-PMID queries (e.g., “Precision Oncology treatment”) showed CosSim = 0.0421 and RCR = 0.89, with FR1 = 0.4726 and FB1 = 0.8571.
- **Interpretation:** With sparse retrievals, semantic relevance dominates—our system reliably summarizes the sole or few matching abstracts, though citation influence plays a reduced role.

#### 6.1.4 Implications

This analysis reveals that query breadth critically affects both retrieval characteristics and ranking effectiveness. Broad queries flood the pipeline with low-impact records, while very narrow queries bypass competition but may underutilize citation data. Moderate queries strike an optimal balance, suggesting that adaptive query guidance (e.g., suggested keyword refinement) could further enhance system performance across diverse information-seeking tasks.

### 6.2 Relation between Ranking Metrics and Summary Quality

#### 6.2.1 High Citation Influence vs. Summarization Performance

- **What we examined:** Queries whose Top-10 Mean RCR exceeded 30 (e.g., “Obesity Prevalence,” “Pulmonary Embolism”).
- **Observations:** “Obesity Prevalence” (Top-10 Mean RCR = 49.27) produced FR1 = 0.5014 and FB1 = 0.8516 (close to dataset averages). “Pulmonary Embolism” (Top-10 Mean RCR = 37.71) yielded the lowest FR1 (0.3922) and one of the lowest FB1 scores (0.8330).
- **Interpretation:** High citation impact does not guarantee strong unigram overlap, as densely formatted epidemiological or clinical abstracts (tables, protocols) limit exact n-gram matching, even though broader context may still be captured semantically.

#### 6.2.2 Semantic Alignment and Summary Quality

- **What we examined:** Queries with Top-10 Mean CosineSim above 0.015 (e.g., “CRISPR gene editing”).
- **Observations:** “CRISPR gene editing” (CosSim = 0.0197) achieved FR1 = 0.5443 and FB1 = 0.8735, among the highest. “Glioblastoma Multiforme” (CosSim = 0.0042) saw below-average FR1 (0.4513) and FB1 (0.8574), despite high citation influence.
- **Interpretation:** Lexical alignment between query keywords and abstract text strongly boosts both ROUGE and BERTScore, underscoring the value of semantic overlap for LLM-based summarization.

#### 6.2.3 Domain-Specific Vocabulary Effects

- **What we examined:** Performance across three domains—Genetics & Molecular Biology, Public Health, and Rare Diseases.
- **Observations:** Genetics queries (e.g., “BRCA1 mutation”) had FR1 = 0.55, FB1 = 0.87. Public Health queries (e.g., “vaccine hesitancy”) showed moderate CosSim (0.010–0.011) but strong FB1 (*σ ≈* 0.85). Rare Disease queries (e.g., “Huntington’s Disease”) returned large PMIDs sets with low median CosSim (*σ ≈* 0.003 ̆ 0.004) and FR1 = 0.50.
- **Interpretation:** Domain jargon alignment matters. Fields with concise, keyword-rich abstracts empower better lexical matches, while heterogeneous or highly technical vocabularies benefit more from semantic metrics.

#### 6.2.4 Implications

This analysis reveals that semantic alignment (CosineSim) often outweighs pure citation influence (RCR) for achieving high summary fidelity. To bolster performance across diverse queries, future work should explore:

- Dynamic ranking weights (e.g., *α* = 0.4*RCR* + 0.6*CosineSim*) based on observed corpus characteristics
- Domain-adapted embedding models for enhanced semantic matching
- User-guided keyword refinement to improve initial lexical alignment
- Hybrid ensembling or re-ranking that tightens n-gram overlap without sacrificing conceptual accuracy

### 6.3 Runtime and Cost Trade-Offs

#### 6.3.1 Effect of Retrieval Size on Latency

- **What we examined:** End-to-end summarization times across queries with varied numbers of PMIDs.
- **Observations:** Sparse queries (1–3 PMIDs, e.g., “wearable health monitors,” “Precision Oncology treatment”) completed in under 24 seconds, with most time spent on dual LLM calls. Large queries (*≥* 200 PMIDs, e.g., “Pulmonary Embolism,” “CRISPR gene editing”) took 25–30 seconds on average, driven by `Entrez.efetch` batching (100 PMIDs per call) and PDF/XML parsing overhead.
- **Interpretation:** Latency remains interactive for niche searches but grows non-linearly with corpus size due to network and parsing delays, suggesting a need for parallelized fetching or local data caching in production environments.

#### 6.3.2 Token-Based Cost Analysis

- **What we examined:** Approximate API token usage and monetary cost per test under a 200-word summary budget.
- **Observations:** Each test used ≈ 1,560 tokens (6 × 260 in max_tokens), costing $0.05–$0.10 per test (i.e., $1–$2 over 20 queries). Scaling to 500-word summaries (≈ 650 in max_tokens) increases costs to $0.20–$0.30 per test ($4–$6 over 20 queries).
- **Interpretation:** While single-user experiments remain inexpensive, costs scale linearly with verbosity—underscoring the value of adaptive word counts, bulk pricing plans, or on-premises inference for large-scale deployments.

#### 6.3.3 Implications

Balancing responsiveness and affordability requires:

- **Adaptive Batching:** Dynamically adjust API batch sizes or parallelize fetch requests to flatten latency curves.
- **Word-Count Tuning:** Tailoring summary lengths per query or user need to minimize token usage.
- **Local Inference:** Exploring open-source LLMs hosted on-premises to eliminate per-token costs.
- **Caching and Previewing:** Storing intermediate representations (e.g., abstracts or embeddings) to reduce redundant processing.

## 7 Discussion

Our personalized literature review system demonstrates significant potential for accelerating scientific sense-making in biomedical domains by effectively addressing the challenge of information overload. The comprehensive evaluation, encompassing both quantitative performance analysis and qualitative human user feedback, reveals key insights into the efficacy of our integrated pipeline and offers clear directions for future advancements.

The *Analysis of Results* illuminated the critical impact of query specificity on retrieval and ranking effectiveness. Our findings indicate that while broad topics can flood the pipeline with numerous low-impact records, and very narrow queries may underutilize citation data, moderately specific queries strike an optimal balance, effectively elevating landmark studies through our composite RCR + Cosine Similarity ranking strategy. This underscores the need for adaptive query guidance mechanisms to optimize information retrieval across diverse user needs.

Furthermore, our analysis revealed that semantic alignment (Cosine Similarity) often outweighs pure citation influence (RCR) in achieving high summary fidelity. This highlights the crucial role of lexical alignment between query keywords and abstract text for LLM-based summarization, especially in fields with concise, keyword-rich abstracts. While high citation impact does not consistently guarantee strong unigram overlap, broader contextual understanding is often still captured. This suggests that enhancing semantic matching through domain-adapted embedding models and user-guided keyword refinement are promising avenues for future improvement.

The *Human User Evaluation* reinforces the system’s practical utility and user satisfaction, with participants consistently rating components highly across navigation, keyword accuracy, paper relevance, and summary quality. The qualitative feedback provides invaluable insights, pointing to the demand for features like expanded corpora, enhanced keyword control, metric transparency, and annotation capabilities. Users particularly valued the inclusion of methodological highlights and author-affiliation context within summaries, affirming the system’s ability to accelerate their appraisal of high-impact literature. While the sample size of 10 users limits broad generalization, the consistent high ratings and thematic feedback strongly indicate the system’s promise and provide concrete directions for UI and underlying retrieval/summarization refinements for broader adoption. Finally, our assessment of runtime and cost trade-offs provides crucial insights for practical deployment. While the system remains interactive for niche searches, latency grows non-linearly with corpus size due indicating the necessity for parallelized fetching or local data caching. Similarly, token-based cost scales linearly with summary verbosity, emphasizing the value of adaptive word counts and potentially exploring local inference for large-scale deployments. These considerations are vital for transitioning the system from experimental use to robust production environments.

Overall, the findings from both quantitative analyses and qualitative user feedback demonstrate that our hybrid ranking and ensemble LLM summarization approach is effective in supporting personalized literature review. The interplay between citation metrics and semantic similarity, combined with dual-model LLM power, offers a powerful paradigm for navigating the ever-growing biomedical literature.

### 7.1 Limitations

Despite the promising results, this study has several limitations. The human user evaluation was conducted with a relatively small sample size (10 participants), which limits the generalizability of the qualitative findings. While the consistency of high ratings is encouraging, larger-scale studies across diverse user groups would provide stronger evidence of usability and effectiveness. Additionally, the system’s current reliance on specific APIs (PubMed, iCite) and fixed ranking weights may limit its adaptability to evolving data sources or user preferences, especially for very broad or very narrow queries where query breadth critically affected retrieval and ranking effectiveness. The identified performance trade-offs in runtime and cost also indicate scalability challenges that need to be addressed for widespread adoption.

### 7.2 Future Work

#### Cross-Domain Adaptation

Our current pipeline leverages MeSH and ACM health ontologies for keyword extraction and ranking. To generalize beyond biomedicine, we will develop a modular ontology plug-in interface, enabling seamless swapping of domain vocabularies (e.g., computer-science thesauri, financial taxonomies, environmental science term banks). Large-scale user studies across these fields will evaluate cross-domain retrieval efficacy and guide iterative refinements.

#### Adaptive Intelligence and Human-in-the-Loop

Building on our two-stage RCR + CosineSim ranking, future work will explore dynamic weighting schemes and reinforcement-learning approaches that update prompt structures and ranking heuristics based on real user feedback. We also plan to prototype a live “keyword edit” interface, letting users refine terms mid-session and see immediate impacts on result sets.

#### Privacy-Preserving & Edge-Friendly Deployment

To reduce network dependence and safeguard sensitive documents, we will implement on-device keyword extraction and local metadata caching. Coupled with lightweight, open-source embedding models, this will enable offline summaries while maintaining performance. We’ll benchmark trade-offs in latency, cost, and security under realistic usage scenarios.

Together, these directions aim to extend our system’s applicability, interactivity, and trustworthiness across diverse information-seeking contexts.

## 8 Conclusion

This paper successfully presents and evaluates a novel, interactive pipeline for personalized biomedical literature review. By integrating ontology-aware keyword extraction, a hybrid RCR + CosineSim ranking approach, and dual-model LLM summarization, our system effectively addresses the challenge of information overload for researchers. The pipeline demonstrates strong computational efficiency, generates high-fidelity summaries validated by both quantitative metrics and positive human user feedback, and offers a cost-effective solution for real-time information interaction. These findings underscore the significant potential of hybrid ranking and ensemble LLM summarization to accelerate scientific sense-making, with broad applicability extending to diverse domains beyond biomedicine.

## 9 Acknowledgments

This work was supported in part by the Department of Computer Science at Purdue University and the School of Science at Indiana University–Indianapolis. We thank the NIH iCite team for providing citation metrics, and the developers of LangChain (2024), OpenAI (2024), and Google Generative AI for making their APIs available. The authors declare no competing financial interests. This research did not receive any specific grant from funding agencies in the public, commercial, or not-for-profit sectors

## 10 Declaration of generative AI and AI-assisted technologies in the writing process

During the preparation of this work the authors used ChatGPT for assistance in initial manuscript drafting and linguistic refinement of certain sections. All generated content was subsequently reviewed, edited, and validated by the authors to ensure accuracy and adherence to scholarly standards. The authors take full responsibility for the content of the published article.

## 11 Data Availability Statement

All primary data used in this study are publicly available. Bibliographic metadata (PMIDs, abstracts) were retrieved via the NCBI Entrez API, and citation metrics were obtained from the NIH iCite portal. Processing scripts and cleaned datasets are available from the corresponding author upon reasonable request.

User study materials (survey instruments and de-identified rating scores) are available from the corresponding author upon reasonable request to protect participant privacy. Any additional materials supporting the findings of this study are likewise available on request.

## References

Devlin, J., Chang, M.-W., Lee, K., & Toutanova, K. (2019). Bert: Pre-training of deep bidirectional transformers for language understanding. Proceedings of NAACL-HLT, 4171–4186. 10.48550/arXiv.1810.04805

Hutchins, B. I., Yuan, X., Anderson, J. M., & Santangelo, G. M. (2016). Relative citation ratio (rcr): A new metric that uses citation rates to measure influence at the article level. PLOS Biology, 14 (9), e1002541. 10.1371/journal.pbio.1002541

LangChain. (2024). Langchain documentation [Available at https://www.langchain.com/].

Lee, J., et al. (2020). Biobert: A pre-trained biomedical language representation model for biomedical text mining. Bioinformatics, 36 (4), 1234–1240. 10.48550/arXiv.1901.08746

Lewis, M., et al. (2020). Bart: Denoising sequence-to-sequence pre-training for natural language generation, translation, and comprehension. Proceedings of ACL, 7871–7880. 10.48550/arXiv.1910.13461

Lin, C.-Y. (2004). Rouge: A package for automatic evaluation of summaries. Text Summarization Branches Out, 74–81. https://aclanthology.org/W04-1013/

Manning, C. D., Raghavan, P., & Schütze, H. (2008). Introduction to information retrieval. Cambridge University Press.

National Institutes of Health. (2024). Icite api documentation [Available at https://icite.od.nih.gov].

OpenAI. (2024). Openai api documentation [Available at https://platform.openai.com/docs/].

Salton, C. G. & Buckley. (1988). Term-weighting approaches in automatic text retrieval. Information Processing & Management, 24 (5), 513–523. 10.1016/0306-4573(88)90021-0

Streamlit Inc. (2019). Streamlit: The fastest way to build data apps in python [Available at https://streamlit.io].

Waltman, L., & van Eck, N. J. (2016). Field-normalized citation impact indicators and the choice of an appropriate counting method. Journal of Informetrics, 10 (2), 752–763. 10.1016/j.joi.2015.08.001

Zhang, T., Kishore, V., Wu, F., Weinberger, K., & Artetxe, A. (2020). Bertscore: Evaluating text generation with bert. 10.48550/arXiv.1904.09675

